# Horizontal gene transfer increases microbiome stability

**DOI:** 10.1101/2022.02.25.481914

**Authors:** K. Z. Coyte, C. Stevenson, C. G. Knight, E. Harrison, J. P. J. Hall, M. A. Brockhurst

## Abstract

Genes encoding resistance to stressors, such as antibiotics or environmental pollutants, are widespread across microbiomes, often encoded on mobile genetic elements. Yet despite their prevalence, the impact of resistance genes and their mobility upon the dynamics of microbial communities remains largely unknown. Here we develop eco-evolutionary theory to explore how resistance genes alter the stability of diverse microbiomes. We show that adding resistance genes to a microbiome typically increases its overall stability, particularly for mobile resistance genes with high transfer rates that efficiently spread resistance throughout the community. However, the impact of resistance genes upon the stability of individual taxa varies depending upon the mobility of the resistance gene and the network of ecological interactions within the community. Non-mobile resistance genes can benefit susceptible taxa in cooperative communities, yet damage those in competitive communities. Moreover, whilst the transfer of mobile resistance genes generally increases the stability of previously susceptible recipient taxa to perturbation, it can, counterintuitively, decrease the stability of the originally resistant donor species. We confirmed these theoretical predictions experimentally using competitive soil microcosm communities. Here the stability of a susceptible microbial community to perturbation was increased by adding mobile resistance genes encoded on conjugative plasmids but was decreased when these same genes were encoded on the chromosome. Together these findings highlight the importance of horizontal gene transfer in driving the eco-evolutionary dynamics of diverse microbiomes.

## Background

Diverse microbial communities colonize virtually every habitat on earth, shaping their abiotic environments and the health of their multicellular hosts^1–3^. Stably maintaining a diverse microbial community is critical for overall microbiome performance, ensuring that the presence of beneficial species or desirable metabolic traits are retained over time^4–7^. In particular, it is crucial that microbial communities can robustly withstand perturbations caused by external stressors, such as environmental pollutants or antibiotics, which may otherwise dramatically reduce overall microbiome abundances and diversity^4,8–10^. Antibiotic-induced changes in community composition have been correlated with a range of adverse health outcomes in host-associated microbiomes^11^, while losses in microbial diversity triggered by heavy-metal and other toxic pollution have been linked to reduced nutrient cycling within environmental microbiomes^12^. Yet despite the importance of withstanding perturbations, the forces shaping the stability of microbial communities remain poorly understood.

Existing theoretical work on microbiome stability has focused primarily on the role of ecological factors, developing mathematical models to disentangle how forces such as microbe-microbe interactions or different classes of stressors influence how microbiomes respond to perturbations^13,14^. However, such models have typically assumed that all species within a given microbiome are equally affected by stressors. Perhaps more importantly, these models typically also assume microbial species remain equally susceptible to stressors over time. In practice, antibiotic or toxin resistance genes are prevalent within microbial communities, often encoded on mobile genetic elements such as plasmids or temperate phages, which can rapidly spread within and between microbial species by horizontal gene transfer (HGT)^15–17^. Therefore, not only are species within microbiomes differentially impacted by stressors, but the rapid spread of mobile genetic elements may dynamically alter the susceptibility of individual microbes to these stressors over short periods of time. These resistance genes and their mobility are highly likely to influence overall microbiome stability, yet exactly how remains unknown.

Here we develop eco-evolutionary theory to examine how the presence and mobility of resistance genes within microbial communities shapes microbiome stability. We then test our key predictions using model soil microbiomes exposed to heavy metal perturbations. In general, our modelling predicts that resistance genes increase overall microbiome stability, with this beneficial effect increasing with increasing gene mobility. However, we also find that a resistance gene can have very different impacts on differing community members, depending upon the precise balance of ecological interactions within a given microbiome, and the mobility of the resistance gene itself. Immobile resistance genes may benefit susceptible species in cooperative communities yet damage those in competitive communities. Meanwhile, though the spread of mobile resistance genes tends to increase overall community stability, it can, counterintuitively, decrease the stability of the originally resistant species. Crucially, our experiments support these key predictions, confirming the beneficial impacts of resistance genes and their mobility on average community properties, and recapitulating the adverse impacts of resistance genes on certain community members. Overall, our work highlights the critical importance of eco-evolutionary dynamics and horizontal gene transfer in shaping complex microbiomes.

## Results and Discussion

### Mathematical model of eco-evolutionary microbiome dynamics

To understand the effect of resistance genes and their mobility on microbial community dynamics we developed a simple and generalizable mathematical model of microbiome dynamics, built around the generalized Lotka-Volterra (gLV) equations (Fig 1A). As in previous work^18–22^, our model assumes the growth of each species within a microbiome is determined by the combination of its own intrinsic growth rate (*r_i_*), its competition with kin (*s_i_*,) and the combination of any interactions each taxon has with other community members (*a_ij_*,). Although simple, these gLV models have been shown to well capture and predict the dynamics of both host-associated and environmental microbial communities^18–22^. Now, we extended this basic model to explicitly incorporate a stressor that inhibits (or kills) susceptible cells and a potentially mobile resistance gene that protects cells encoding it, but at the cost of a reduced intrinsic growth rate. Using this model, we could simulate the individual species abundances of any given microbiome over time. More specifically, we could simulate the scenario in which a microbiome equilibrates, but is then briefly exposed to an external stressor such as a heavy metal or an antibiotic perturbation (Fig 1B). By measuring the average change in species abundances during this perturbation we could therefore quantify the stability of that microbiome.

**Figure 1.**
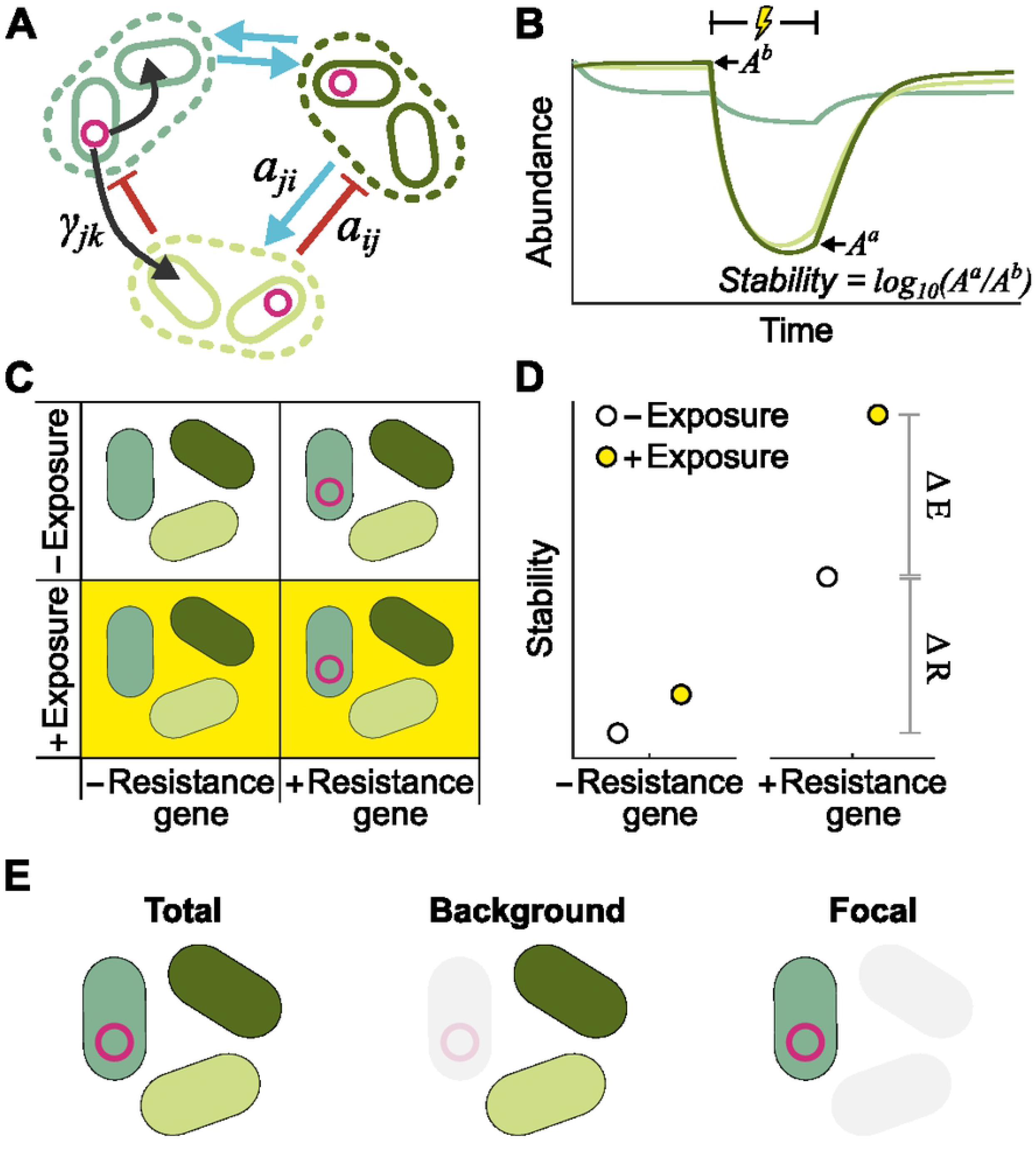
Mathematical modelling captures eco-evolutionary dynamics of microbial communities. **A.** Schematic illustrating our mathematical model; each species (dashed line) is composed of two populations, with and without a resistant gene. Species impact one another’s growth through ecological interactions such as competition or competition (blue and red arrows), while horizontal gene transfer enables resistance genes to spread within and between species (black arrows). **B.** Schematic illustrating representative microbiome dynamics, capturing species dynamics before, during, and after an external perturbation (lightning bolt). We calculate each species’s abundance immediately before, *A^b^*, and after, *A^a^*, the perturbation period, then define stability as the average logged fold change for each species (mean log_10_(A^a^/A^b^)) **C.** Schematic illustrating our four modelling scenarios: communities with and without resistance genes, and with and without prior exposure to low-level stressors. **D.** Comparing the four scenarios allows us to calculate the change in microbiome stability that results from the introduction of a resistance gene (ΔR), and the change in stability that results from prior exposure to weak low-level selection (ΔE). **E.** To disentangle the impact of resistance and selection on different taxa we calculate ΔR and ΔE for the total community, the background community only and the focal species alone.

Having established this basic model, we used it to explore the impact of resistance genes on the stability of microbiome communities. Specifically, we generated a series of diverse multispecies microbial communities, then quantified the stability of each of these microbiomes under four distinct scenarios (Fig 1C). First, we simulated microbiome dynamics when all species were susceptible to the stressor, and then again when a randomly chosen focal species carried a resistance gene. Next, we repeated this process, but allowed each microbiome to initially equilibrate in the presence of a low level of the stressor, for example, capturing prior exposure to subinhibitory levels of antibiotics or pollutants. This process allows us to define two metrics: the change in microbiome stability resulting from the introduction of a resistance gene, ΔR, and the change in microbiome stability resulting from prior exposure to low-level selection, ΔE (Fig 1D). For each community, we calculated ΔR and ΔE for the whole microbiome (Fig 1E), the focal species only (i.e., the species that originally carried the resistance gene), and the background community only (i.e., all species except the focal species). Using this modelling framework, we could then explore in depth the impact of resistance genes on microbiome stability.

### Mobile resistance genes increase stability of non-interacting communities

We began by exploring how the presence and mobility of resistance genes influenced the stability of microbiomes in the absence of between-species ecological interactions (that is, all *a_ij_* = 0). To do so, we generated a set of communities with and without resistance genes. We then systematically varied the ability of these resistance genes to transfer within and between species (Fig 2A), capturing all degrees of gene mobility from immobile (e.g., a chromosomally encoded resistance gene) to highly mobile (e.g., a resistance gene encoded by a highly conjugative and promiscuous plasmid).

**Figure 2.**
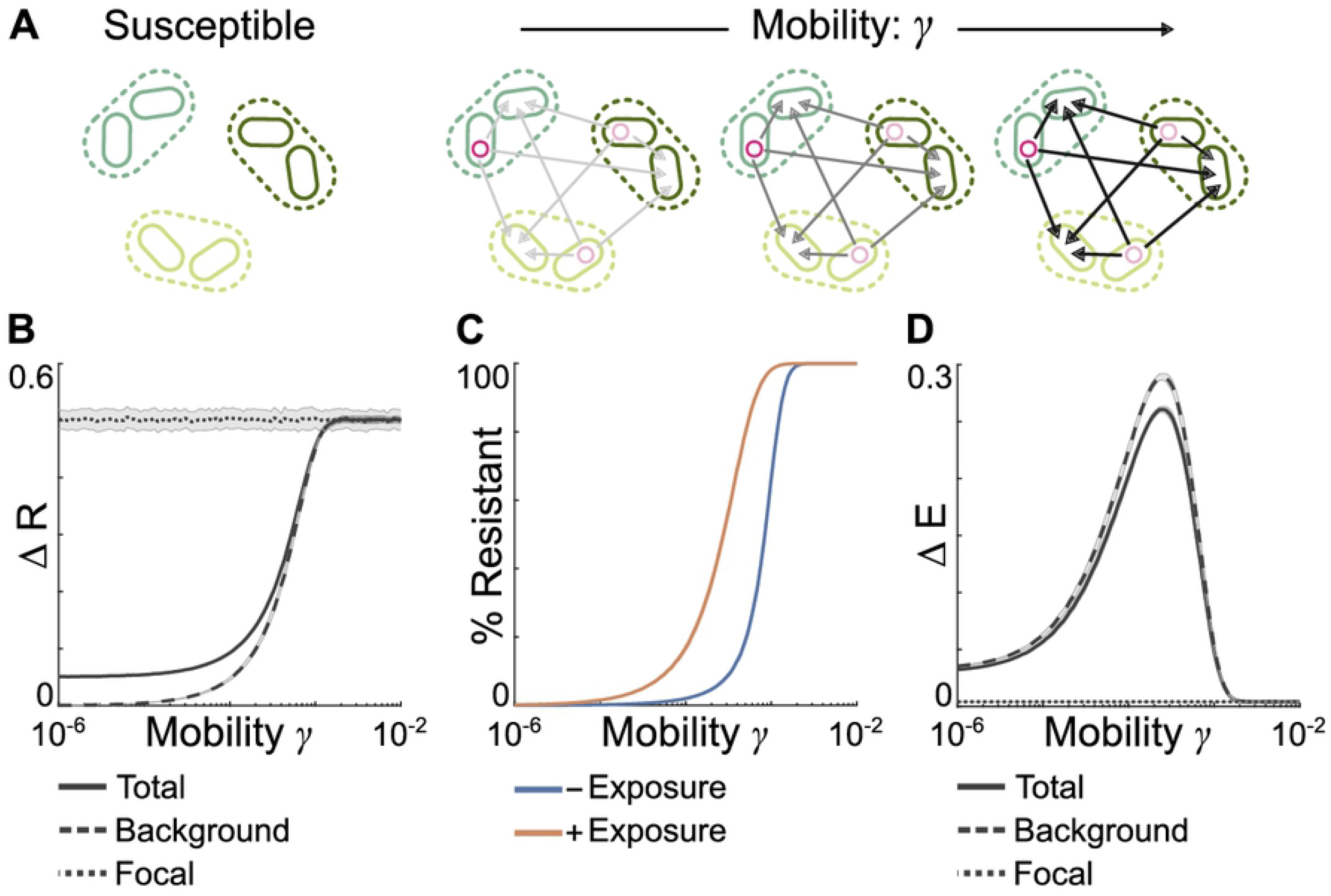
In simple communities mobile resistance genes increase microbiome stability. **A.** Schematic illustrating our modelling approach. We generate a set of simple microbiomes without between species interactions, then for each community we simulate the introduction of a series of resistance genes with increasing mobility. **B.** ΔR, the change in stability resulting from the introduction of a resistance gene, for the community as a whole (solid line), the background community (dashed line) and the focal species (dotted line). Any resistance gene significantly increases overall community stability, but only highly mobile resistance genes substantially increase the stability of background species. **C.** Plasmid frequency within the background community with (orange) and without (blue) prior exposure to low-level stressors. Prior exposure increases plasmid frequency, with this increase greatest for plasmids with intermediate mobility. **D.** ΔE, the change in stability resulting from prior exposure to low stressor levels, for the community as a whole (solid line), the background community (dashed line) and the focal species (dotted lined). Prior low-level selection increases the stability of both the community as a whole and background species, with this effect greatest for communities with intermediate mobility plasmids. Throughout lines and error bars represent mean and standard deviation over 100 independent, 10-species communities, with model parameters given in Table 1.

Introducing any resistance gene increased overall microbiome stability, decreasing the average drop in community abundances following the onset of the perturbation (Fig 2B, ΔR > 0). However, examining background and focal species separately revealed that in most cases this increase was driven solely by the increased stability of the focal species. That is, the presence of a resistance gene in the focal species drove up the average stability of the community as a whole, but in most cases the stability of background species remained effectively unchanged (Fig 2B). To substantially increase the stability of the background microbiome, we found that the novel resistance gene must be highly mobile. This was because, prior to the perturbation, the resistance gene did not confer any benefit, and thus could only spread into the background community when its transfer rate exceeded its rate of decline caused by negative selection against its cost. However, low-mobility resistance genes could increase background species stability provided additional forces enhanced the spread of resistance prior to any perturbation. For example, prior exposure to weak stressor selection introduced a weak benefit to harboring the resistance gene prior to the perturbation, enabling it to spread into the background community even at lower rates of gene mobility (Fig 2C). As a consequence, prior exposure substantially increased the stability of background species, even in communities harboring low-mobility resistance genes (Fig 2D).

### Interspecies interactions modulate the impact of resistance genes on microbiome stability

Having established these baseline properties of the system, we next examined how the impact of resistance genes upon community stability is modulated by interspecies interactions. Specifically, we allowed individual species within our simulated communities to interact with one another in a variety of different ways, ranging from competition and ammensalism (-/- and -/0 interactions respectively), through exploitation (+/-), to cooperation and commensalism (+/+ and +/0). We then generated a range of different microbial communities, systematically varying the proportion of each interaction type (Fig 3A). As previously, we then simulated the introduction of resistance genes into these communities, also systematically changing the mobility of the resistance gene.

**Figure 3.**
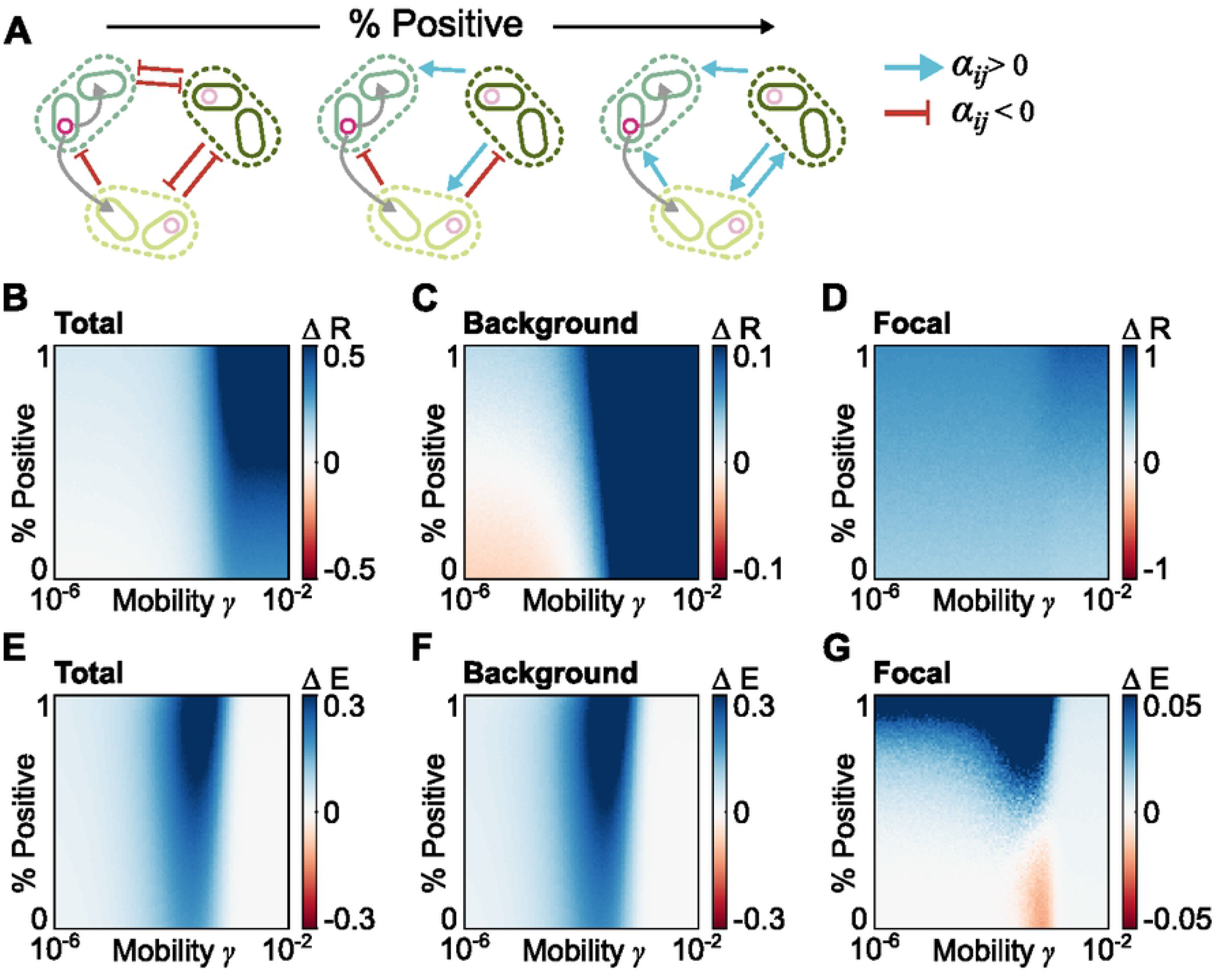
Interspecies interactions modulate the effect of resistance genes. **A.** Schematic illustrating our modelling approach. We generated a series of diverse microbiomes, systematically increasing the frequency with which microbes facilitate one another’s growth. **B-D.** Average ΔR, the change in stability resulting from the introduction of a resistance gene, under varying community types and plasmid transfer rates, shown for the whole microbiome (B), the background community (C) and the focal species alone (D). **E-G.** Average ΔE, the change in stability resulting from prior exposure to low stressor levels, under varying community types and plasmid transfer rates, shown for the whole microbiome (E), the background community (F) and the focal species alone (G). Throughout patch color represents mean ΔR or ΔE over 100 independent, 10-species communities, across a range of 101 Positivity and *γ* values. Other model parameters given in Table 1.

As in non-interacting communities, introducing any resistance gene typically increased overall community stability, and this increase was higher for more mobile resistance genes (Fig 3B, S1-5). However, this beneficial effect of resistance genes varied with interaction type and was far stronger in microbiomes with a high proportion of cooperative interactions. In these cooperative communities, individual species benefited both directly from acquiring resistance genes, and indirectly from their cooperative partners acquiring resistance, which in turn helped to buffer the negative impact of any perturbation. The principal effect of prior weak stressor exposure was once again to reduce the level of gene mobility required for resistance genes to spread within the community. And, as a consequence, this prior stressor exposure increased the stabilizing effect of mobile resistance genes on overall community stability, across all interaction types (Fig 3E, S1-5, although, notably this effect was strongest for resistance genes with intermediate mobility).

While mobile resistance genes were beneficial for overall microbiome stability, examining focal and background species separately revealed surprisingly different dynamics – with background and focal species showing markedly different responses to resistance genes depending upon the precise manner in which species were interacting and the mobility of the resistance gene (Fig 3C, D, S1-5). In cooperative microbiomes, background species benefited from the introduction of a resistance gene regardless of its mobility. This occurred because, by promoting the survival of species with whom a susceptible species cooperates, resistance genes aid recovery of the susceptible species regardless of whether they have access to the resistance gene through HGT. However, in competitive communities highly mobile resistance genes increased background community stability while low-mobility genes *reduced* background community stability (Fig 3C). That is, in highly competitive communities most species were less stable when another member of the community harbored an immobile or low mobility resistance gene than when all species were susceptible. What drove this phenomenon? In fully susceptible competitive communities, during a perturbation every species experienced a reduction in their net growth rate, typically resulting in a decrease in their overall abundance. However, as a consequence each species also benefited from some competitive release – that is, the negative impact of competitors was reduced as these competitors also decreased in abundance. In contrast, if one species acquired an immobile or low-mobility resistance gene then this focal species remained at a high density during the perturbation – and as such, susceptible species not only suffered from the stressor-mediated inhibition, but also from continued strong competitive inhibition by the focal species.

This impact of competitive release also modulated the impact of prior selection – again with very different impacts on background and focal species. Background species benefited from prior exposure to low-level stressors regardless of community context (Fig 3F), as this initial selection promoted the spread of mobile resistance into the community. Moreover, this spread of mobile resistance also stabilized cooperative focal species, as these species benefited from their cooperative partners acquiring resistance genes and thus remaining at high densities during perturbations (Fig 3G). However, in certain competitive communities prior selection could, counterintuitively, slightly reduce the stability of the focal species (ΔE<0, Fig 3G) because the spread of mobile resistance into background species meant that the focal species no longer benefited from any competitive release during perturbations. Altogether, our results suggest the introduction of a mobile resistance gene can have a wide variety of effects, with the exact impact depending upon which species are being examined, how they interact with one another, and the mobility of the resistance gene.

### Experimental microbial communities reproduce theoretical predictions

To test our predictions about the impact of resistance genes and their mobility on community stability we developed an experimental model microbiome system. Specifically, we generated model microbiomes by inoculating sterile potting soil microcosms with 96 soil bacterial isolates (our background community) and one focal species *Pseudomonas fluorescens* SBW25. In each microcosm this focal species was either susceptible to mercury (Hg^S^), or carried a mercury resistance operon, encoded either on the chromosome or on one of two conjugative plasmids, pQBR103 and pQBR57. Whereas the chromosomally encoded resistance is non-mobile, the conjugative plasmids can transfer mercury resistance to other species^23–25^. Having assembled these communities, we allowed them to equilibrate for ten serial transfers (40 days) either with or without weak mercury selection (Fig 4A). After equilibrating, communities were perturbed with a pulse of high concentration mercury, then propagated without mercury for two additional serial transfers, mirroring the mode of perturbation used in our modelling framework. We determined the composition of bacterial communities by 16S rDNA gene amplicon sequencing before and after the perturbation, and estimated the abundances of the total community by colony counts. As in our models, we then quantified community stability as the change in species abundances between the pre- and post-perturbation samples.

**Figure 4.**
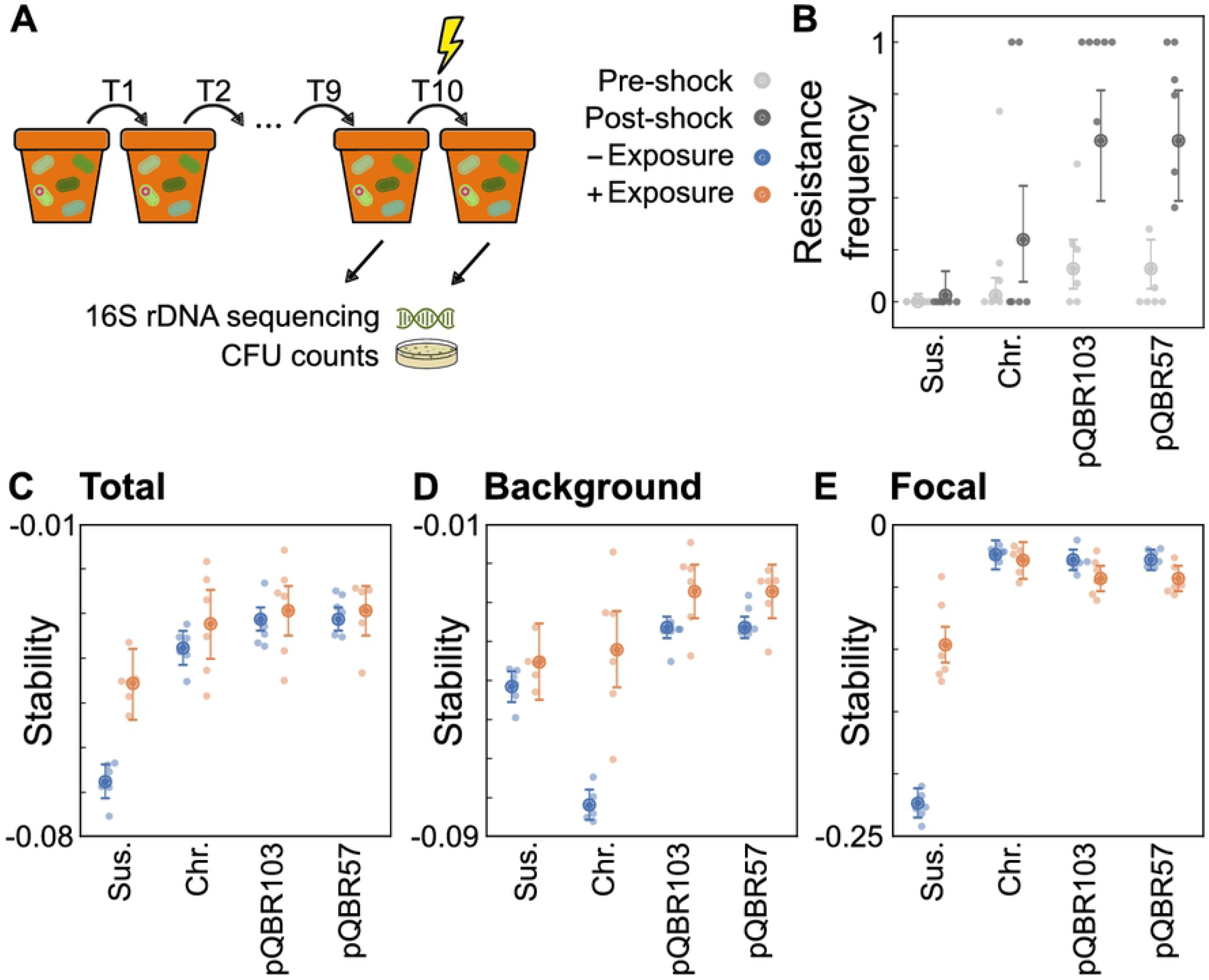
Experimental microbiomes recapitulate the predicted impact of resistance genes on microbiome stability. **A**. Schematic illustrating our experimental system. Here soil microcosms are seeded with microbial communities, allowed to equilibrate with or without low-level mercury, then perturbed with a high-level mercury pulse. **B.** Frequency of resistance within the background population before (light grey) and after (dark grey) the high-level mercury pulse, calculated for each experimental condition (fully susceptible, Sus., and Chromosomal, Chr. or plasmid-carried resistance, pQBR103, pQBR57) **C.** Total microbiome stability, as measured by log10(Fold Change) in species abundances following perturbation, such that more negative values indicate a less stable community. **D.** Stability of the background community individually. **E.** Stability of the focal species alone. Throughout, orange and blue dots indicate community stability with and without prior mercury exposure respectively, with each condition (resistance x prior selection) containing n=6 independent samples.

After equilibrating, each microbiome contained between three and five taxa at appreciable abundances, namely: OTU_107 (the focal species, *P. fluorescens* SBW25), OTU_3 (*Pseudomonas sp.*), OTU_14 (*Pseudomonas umsongensis*), OTU_9 (*Bacillus megaterium*), and OTU_19 (*Bacillus simplex*). Although the exact ecological interactions occurring between each of these taxa are not known, there is good evidence for exploitative and interference competition occurring between Pseudomonas species^26,27^, and between members of the Bacillus and Pseudomonas genera^28–31^, suggesting this is likely to have been a competitive community. As such, our model predicts that an introduced resistance gene would be likely to increase overall microbiome stability, but that background species would only benefit from plasmid-encoded resistance, and might suffer from the introduction of chromosomally encoded genes. Moreover, our modelling also predicts that prior selection might, counterintuitively, slightly decrease the stability of the focal species when it harbored plasmid-encoded mercury resistance, due to transfer of mobile resistance limiting the benefits of competitive release.

As predicted, the presence of any resistance gene in the focal species SBW25 significantly increased overall microbiome stability, strongly reducing the decline in abundance caused by the mercury pulse (Fig 4C, effect of resistance = 0.030 [0.025 – 0.035, 95%CI], Bayesian linear model BLM1, see SI). Similarly, as predicted, there was an overall positive effect on stability of prior exposure to weak mercury selection (effect of exposure = 0.022 [0.013 – 0.031, 95%CI], BLM1). We also observed a positive impact of gene mobility on overall microbiome stability, however, this effect was small (estimated effect of mobility on stability, given resistance = 0.0065 [0.0017 – 0.011, 95% CI], BLM1), and did not change significantly with prior exposure (estimated interaction effect between exposure and mobility, given resistance = −0.0035 [−0.014 – 0.0075 95% CI], BLM1) – reflecting that increases in stability were dominated by the behavior of the focal species.

In contrast, and as predicted by our mathematical modelling, calculating the stability of background and focal species members separately revealed markedly different behavior between species, and a far stronger effect of resistance gene mobility (Fig 4D). In communities without prior selection, plasmid-encoded mobile resistance genes increased background community stability relative to fully susceptible communities (estimated effect of resistance on stability, given mobility = 0.015 [0.010 – 0.020], Bayesian linear model BLM2, see SI). And, consistent with this increased stability being driven by plasmid transfer into the background community, in these communities we also observed significantly higher levels of mercury resistance in the background species following the perturbation when plasmids were present (Fig 4D, effect of mobility on resistance frequency, given resistance = 0.21 [0.064 – 0.35] Bayesian linear model BLM3). In contrast, however, adding immobile chromosomal resistance genes strongly *reduced* background community stability (Fig. 4D, effect of chromosomal resistance = −0.030 [−0.036 – −0.025, 95% CI], BLM 2). That is, in microbiomes harboring immobile resistance genes, background species were more strongly perturbed by the mercury pulse than in microbiomes entirely lacking resistance genes. Moreover, immobile resistance genes caused an even greater drop in background species stability when compared to communities harboring mobile resistance genes (effect of mobility on stability, given resistance = 0.046 [0.041 – 0.050, 95% CI], BLM2). Together, these results support our predictions that resistance genes can have markedly different impacts on background community members, depending on their mobility.

Finally, we observed a striking difference in the response of the background and focal taxa to prior mercury exposure. Specifically, when resistance was present within the community, prior exposure increased the stability of the background species (effect of exposure given resistance = 0.034 [0.019 – 0.048, 95% CI], BLM 2) yet decreased the stability of the focal species (interaction effect of focal community with exposure, given resistance = −0.17 [−0.19 – −0.14, 95% CI], BLM2). And, notably, this decreased stability was unique to resistant focal species, with prior exposure increasing the stability of the focal species if it was sensitive (interaction effect between focal community and exposure = 0.12 [0.10 – 0.14, 95% CI], BLM2). This suggests that, as predicted by our models, the increased stability of the background species might indeed be reducing the competitive release experienced by the focal species during perturbations, and thus decreasing the overall stability of the focal species following prior exposure (cf. red region in Fig 3G). Collectively, these experimental findings support our theoretical predictions relating to the effects of resistance genes and their mobility upon the stability of competition dominated microbiomes.

## Conclusions

Genes conferring resistance to stressors such as antibiotics, toxins or pollutants are widespread within microbial communities, often encoded on mobile genetic elements such as plasmids or temperate phages^15–17^. While the consequences of resistance – particularly to antibiotics – for human health have been widely studied^32^, the impact that resistance genes have on the structure and stability of microbial communities remains poorly understood. Here we combine novel theory and experiments to disentangle the diverse ways in which resistance genes influence the stability of microbiomes. Our work suggests that resistance genes typically increase overall community stability, particularly when encoded on highly mobile genetic elements. However, exactly how these genes influence microbiome properties depends upon the precise interplay between the properties of the gene and of the underlying microbiome. The same gene may have directly opposing effects upon microbiome stability, depending upon its mobility, or how individual species interact with one another. Moreover, not all species are affected equally and, particularly in highly competitive microbiomes, the presence and transfer of resistance genes may benefit some species yet be detrimental to others.

The considerable variability in the effects of resistance genes within microbiomes introduces an interesting set of potential conflicts between genes, their bacterial hosts, and the broader ecosystem. For example, the spread of antimicrobial resistance poses a dangerous threat to public health^33^. Yet within a given microbiome, the spread of resistance into susceptible taxa may offer important ecological benefits, improving overall microbiome stability and protecting susceptible community members from out-competition by resistant competitors. Increasing community stability through the spread of mobile resistance genes could also enable the maintenance of important ecosystem services within vulnerable microbial communities. Similarly, transfer of a resistance gene into susceptible taxa may be advantageous for the fitness of the individual gene, increasing its frequency within the community. However, in certain microbiomes spread of the resistance gene may be costly for the original host – reducing its advantage over otherwise susceptible competitors^34^. Together, these conflicts reveal the important and sometimes complex effects mobile genetic elements can have upon microbial communities. Moreover, they also underline the importance of considering exactly how properties such as microbiome stability are quantified. Relying solely on coarse, whole-community metrics such as overall community abundances or diversity may mask striking differences between individual community members, and risks obscuring important eco-evolutionary dynamics.

To identify broad patterns in the impact of mobile resistance genes upon microbiomes here we used relatively simple ecological models. The advantage of these simple models is that we can analyze large numbers of microbiomes in high-throughput. However, as a consequence, there are ecological and evolutionary features that we have not explicitly accounted for. For example, previous studies have suggested that plasmid transfer is more likely between closer phylogenetic relatives, due to constraints on plasmid host-range or spatial structuring^35^. Indeed, in our experiments plasmid-mediated transfer of Hg^R^ from *P. fluorescens* SBW25 appeared to be limited to congenerics — with more distantly related members of the community such as *B. megaterium* apparently unable to gain the resistance genes, suggesting they were unable to acquire or maintain either plasmid. Similarly, our model does not explicitly include *de novo* mutations or phenotypic changes that may modulate how individual taxa respond to stressors without requiring the acquisition of resistance genes from other community members. Indeed, in our experiments we observed an increase in the stability of background species after prior exposure to mercury within the chromosomally-encoded resistance treatment (Fig. 4D), suggesting mechanisms other than plasmid transfer may have driven this increased stability^36^. Each of these phenomena has the potential to further modulate the impact of horizontal gene transfer upon microbiome stability, and disentangling exactly how will be critical if we wish to fully understand and ultimately predict microbiome dynamics.

## Materials and Methods

### Underlying microbiome model

In line with previous work^18,19^, we model each microbiome as a network, where each node represents a species and each edge captures the interaction between them. However, now we extend this model to include two populations for each species, one with plasmids and one without^25^. For each species *i*, plasmid-free cells grow at a rate *r_i_* and plasmid-bearing cells grow at a rate *r_i_ - c*, where c indicates the cost of plasmid carriage. Plasmids can transfer within and between species, with the per cell rate of plasmid transfer from species *j* into species *i* defined as 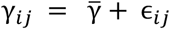, where 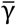 is the average plasmid transfer rate, and ϵ*_ij_* is a noise term drawn from a Normal distribution to introduce variability in plasmid transfer rates between species (note, in instances where γ*_ij_* < 0 we set γ*_ij_* = 0). Finally, we introduce an inhibition term -β*D*, where D defines the level of antibiotic or toxin in the environment, and β the susceptibility of the population in question. More specifically, we set β*^s^* = 1 for the susceptible population, and β*^r^* = 0.1 for the resistant population. Together, this enables us to define the growth rate of the plasmid-negative, *X^s^*, and plasmid-positive, *X^r^*, populations of a given species as,

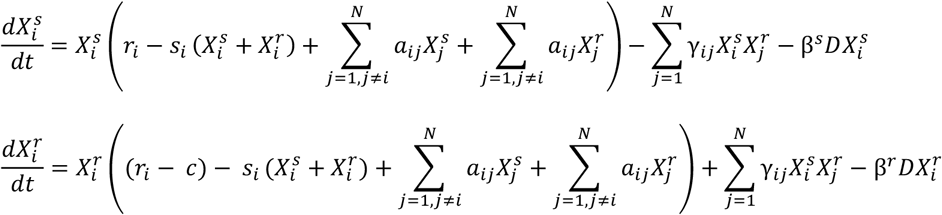

with equivalent expressions defining the dynamics of each other species *j* = 2:*N* within the community. Notably, this model can also be extended to incorporate the phenomenon whereby plasmids can be lost during segregation at a rate δ, however, this does not qualitatively alter the results (Fig S5).

Having established this new model, we generated a series of microbiomes, each composed of *N*=10 species. Within any given microbiome, each species i interacts with species j with probability *C*. To investigate how interspecies interactions modulate the effect of resistance genes, we systematically alter the proportion of individual interaction terms, *a_ij_*, that are positive, P_m_, such that when P_m_ = 0 the community is purely competitive, when P_m_ = 0.5 the community contains a mixture of all interaction types, from competitive and ammensal, through exploitation, to cooperative and commensal, and when P_m_ = 1 the community is purely cooperative. Finally, we choose the magnitude of each *a_ij_* is from a half-normal distribution with standard deviation σ = 0.05, and set the intrinsic growth rates *r_ij_* such that the community has a linearly asymptotically stable equilibrium 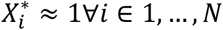 in the absence of any resistance gene. Importantly, we find similar results when varying each of these key microbiome parameters (Figs S3-5).

### Quantifying microbiome stability

We calculate the stability of any given community by simulating its dynamics in response to a perturbation. Specifically, we allow the community to equilibrate, then briefly perturb it by setting the stressor level within the environment to D = 0.1 for t = 25 time units. We then measure the difference in abundance of each species *i* before, 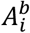, and after, 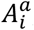, this perturbation. We define the stability of each individual species *i* based on their fold change in response to the perturbation, such that 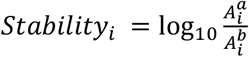, then define the stability of the whole community (or background community), as the mean of *Stability_i_* across all species (or across only the background species). To explore the impact of resistance genes on stability for each community we perform these simulations when all species are susceptible to the stressor, and when one randomly chosen species harbors a mobile resistance gene. To explore the impact of low-level selection on stability, we perform these simulations when the community initially equilibrates in a stressor-free environment, and when the community equilibrates in the presence of a low-level of the stressor (setting D = 0.01 during the equilibration period), with all simulations performed in MATLAB R2020a.

### Strains and culture conditions

To test the accuracy of our predictions, we assembled experimental model microbiomes, composed of a defined background community augmented with a predetermined focal species. For our focal species we used *Pseudomonas fluorescens* SBW25 labelled with a gentamicin resistance marker using the mini-Tn7 transposon system^25,36–38^. More specifically, we generated independent *P. fluorescens* strains that were either susceptible to mercury, or harbored harboured a mercury resistance gene, Hg^R^, on either the chromosome, the conjugative plasmid pQBR57^38,39^, or the conjugative plasmid pQBR103^39^. Individual colonies of each strain (one for each replicate) were isolated on selective KB agar and grown overnight at 28 degrees in 6ml KB broth (10 g glycerol, 20g proteose peptone no. 3, 1.5g K_2_HPO_4_•3H_2_O, 1.5g MgSO_4_•7H_2_O, per litre^40^) in a shaking incubator.

To generate our background community, we plated supernatant from non-autoclaved John Innes No.2 potting soil on nutrient agar plates, which were then grown for 48 hours in a 28 degree centigrade incubator. Following this incubation period, we randomly selected 96 colonies, which we screened against Hg^2+^ and Gm to ensure no community members already harbored phenotypic resistance to either our stressor or selective marker. Each of these bacterial strains were grown separately overnight in 6ml KB broth in a shaking incubator and then washed and mixed at an equal volume to make the background community. To generate each final microbiome community, we resuspended an equal volume of the background community and an overnight culture of *P. fluorescens* in M9 buffer. We then diluted this suspension by 1:10 with M9 buffer, and used 100μl to initiate each experimental replicate. Populations were cultured in sterile soil microcosms consisting of 10 grams twice autoclaved John Innes No.2 potting soil supplemented with 900μl of sterile H_2_O and maintained at 28 degrees at 80% relative humidity.

### Community perturbation experiment

We established twelve replicate communities per SBW25 genotype: SBW25, SBW25 with chromosomal Hg^R^, SBW25 carrying Hg^R^ encoded on pQBR57, SBW25 carrying Hg^R^ encoded on pQBR103. These were propagated by serial transfer of 1% of the community every 4 days into fresh soil microcosms for twelve transfers. In half of these lines communities were augmented with mercuric chloride (at 16 μg/g HgCl_2_) from transfer 2 onwards, allowing us to capture communities with and without prior stressor exposure. At transfer 10 all communities were perturbed by exposure to 128 μg/g HgCl_2_ and then propagated for a further serial transfer at 0 μg/g HgCl_2_. Samples of each community were cryogenically stored at each serial transfer in 20% glycerol. Throughout the experiment, we determined the population density of SBW25 by diluting and plating samples onto KB agar supplemented with 6 μg/ml gentamicin, and the abundance of the entire community by plating onto nutrient agar. In addition, we determined the frequency of the Hg^R^ phenotype in the community as a whole by plating onto nutrient agar supplemented with 20μM HgCl_2_, while we determined the frequency of Hg^R^ resistance in the focal strain by plating onto nutrient agar supplemented with 20μM HgCl2 and 6 μg/ml gentamicin.

### 16S rRNA gene sequencing and analysis

To quantify community composition we extracted whole community DNA samples from the thawed stocks stored on days 10 and 11 (i.e. before and after the mercury shock, Fig 4a). Specifically, we extracted DNA using QIAGEN DNeasy PowerSoil kits, following manufacturer instructions with the exception that stocks were initially spun down and re-suspended in 1x M9 to remove glycerol. DNA concentrations were assessed using Qubit fluorometer 3.0 (Thermo Fisher Scientific) and diluted to 20ng μl^-1^ where possible before samples were sent for downstream PCR amplification of the v4 region of 16s rRNA gene and sequencing on the Illumina MiSeq platform. The raw forward and reverse reads were merged and processed using QIIME1^41^. Reads were stripped of their primer and barcoding sequences using Cutadapt^42^ and untrimmed reads were discarded. Reads were truncated at 254 bp (size of the amplicon). Reads were then dereplicated using Vsearch^43^ and clustered into operational taxonomic units (OTUs) using Usearch^44^ with 97% similarity. OTUs were then filtered based on OTUs which appeared in the positive control (14 OTUs in total)._Putative taxonomic identification of the 14 OTU’s was performed using BLAST^45^ to align the OTU sequence data to the NCBI nucleotide database, listed in SI Table 2. Total abundances of the focal species and the background community were determined by multiplying the relative abundances of each with the total community abundances calculated by CFU counts.

### Statistics

Differences in microbiome stability and resistance frequency were estimated using three Bayesian linear models, accounting for the experimental structure (where relevant, the presence of resistance, its mobility, foreground versus background communities, prior exposure to weak mercury selection and timing of measurement relative to mercury shock), non-homogeneity of variances and, where appropriate, non-normality of residuals, using broad priors. These models were fitted with the brms package^46^ (version 2.16.3) which uses STAN via the R language^47^ (version 4.1.2). Four MCMC chains were used, each of 4,000 iterations, where the first 2,000 were discarded as warm-up, resulting in 8,000 draws from the posterior distribution. Convergence was checked visually using plots of the draws and via the R-hat value^48^, which will equal 1 at convergence and was 1.0 for all parameter estimates reported in the main text and Supplemental Information. All values are reported as a mean with 95% credibility interval (CI). Details of model structures and estimated parameters for each of the three models (Supplemental Tables 3-5) are given in the Supplemental Information.

## Acknowledgements

We thank D.R. Gifford and W.P.J. Smith for helpful discussions. K.Z.C. was funded by a University of Manchester Presidential Fellowship. C.S. was supported by an ACCE DTP NERC PhD studentship. M.A.B., J.P.J.H. and E.H. were funded by grants BB/R014884/1 and NE/R008825/1.

